# Advancing Antibiotic Resistance Classification with Deep Learning Using Protein Sequence and Structure

**DOI:** 10.1101/2022.10.06.511103

**Authors:** Aymen Qabel, Sofiane Ennadir, Giannis Nikolentzos, Johannes F. Lutzeyer, Michail Chatzianastasis, Henrik Boström, Michalis Vazirgiannis

## Abstract

**Background:** Antibiotic resistance is a major global health concern, as bacteria can develop immunity to drugs rendering them ineffective. To address this problem, it is crucial to identify and classify the genes that are responsible for antibiotic resistance, i. e., antibiotic resistant genes (ARGs). Previous methods for gene classification have mainly focused on the sequence of proteins and have ignored their structure. Recently, the AlphaFold model has made significant progress in predicting the 3D structure of proteins. Since the sequence and the structure can complement each other, having access to both of them can allow machine learning models to more accurately classify novel ARGs. In this paper, we develop two deep learning models to classify novel Antibiotic Resistant Genes (ARGs) using information from both protein sequence and structure. The first architecture is a graph neural network (GNN) model equipped with node features derived from a large language model, while the second model is a convolutional neural network (CNN) applied to images extracted from the protein structures.

**Results:** Evaluation of the proposed models on a standard benchmark dataset of ARGs over 18 antibiotic resistance categories demonstrates that both models can achieve high accuracy in classifying ARGs (> 73%). The GNN model outperformed state-of-the-art methods and provided rich protein embeddings that could be also utilized in other tasks involving proteins, while the CNN model achieved competitive performance. With larger datasets, it is expected that the performance would further increase due to the nature of the underlying neural networks.

**Conclusions:** The proposed deep learning methods offer a more accurate approach for antibiotic resistance classification and hold significant potential for improving our understanding of the mechanisms underlying antibiotic resistance.

## Background

Humans and bacteria have a long-standing symbiotic relationship which dates back to the dawn of humanity. Some bacteria are very useful for humans since they play a key role in some functions. However, there also exist bacteria which are responsible for certain human diseases such as meningitis and tuberculosis. Since 1928 when Alexander Fleming discovered penicillin, antibiotics have served as the main weapon to treat and prevent bacterial infection. Antibiotics have thus contributed to an increase in life expectancy and to the improvement of healthcare systems since surgeries and other types of operations can now be carried out with less risk for severe consequences from bacterial infections [1, 2].

However, due to the overuse and misuse of antibiotics, bacteria have become resistant and no longer respond to these drugs, a phenomenon known as antibiotic resistance. These resistant bacteria cause infections that are harder to treat resulting in higher medical costs and increased mortality. In addition to the overuse of antibiotics, the rise of antibiotic resistance is also due to the inability of the pharmaceutical industry to develop new drugs, which is due to challenging regulatory requirements among others. Antibiotic resistance is estimated to be responsible for over 33,000 deaths across Europe in 2015 [3], and if not addressed, it is expected that by 2050, around 10 million people globally could be at risk every year, with an economic cost exceeding $100 trillion [4].

Thus, it becomes clear that there is an urgent need for actions to prevent the antibiotic resistance crisis. Some examples of such actions include reducing the unnecessary use of antibiotics and using optimal treatments to cure infections. Antibiotic resistance usually occurs as the result of genetic mutations. However, bacteria can also become resistant to antibiotics by the acquisition of specific genes through horizontal gene transfer [5]. Therefore, finding the antibiotic resistant genes (ARGs) along with the antibiotic class they are resistant to is of paramount importance for the development of targeted treatment, but also for the prevention of infections.

To that end, the research community has resorted to computational methods, among others, which can compare sampled genome sequences against sequences that exist in reference databases. These methods typically employ well-established sequence alignment algorithms such as BLAST [6] and DIAMOND [7]. Most of these methods use existing microbial resistance databases along with a “best hit” approach to predict whether a sequence is indeed an ARG.

While the aforementioned approaches can accurately recognize known or highly conserved sequences of ARGs, they may fail to detect novel ARGs or those that exhibit low levels of similarity to known ARGs. This results in a significant amount of false negative samples, i.e., several ARGs are classified as not being ARGs [8, 9, 10]. Recently, some deep learning approaches have been proposed to deal with the above limitations. For example, DeepARG [8] generates features that capture the distances of a given sequence to known ARGs and passes them onto a deep learning architecture. This method was shown to improve the classification performance and to decrease the false negative rate. Hamid and Friedberg investigated the use of transfer learning in the ARG classification setting [11]. They first constructed a new dataset, called COALA (COllection of ALl Antibiotic resistance gene databases), by combining 15 different datasets, and then pre-trained an LSTM-based language model using the ULMFit (Universal Language Model Fine-tuning) method [12] on 734,848 bacterial proteins sampled from the UniRef50 database [13]. The model was subsequently trained on the newly created COALA dataset where it outperformed alignment-based methods. Another recent method, ARG-SHINE [14], integrates multiple techniques in an ensemble. More specifically, it aggregates the results of a convolutional neural network (CNN) applied to the sequences, a protein functional-based logistic regression model, and a BLAST-based model. The ARG-SHINE ensemble was found to outperform the aforementioned approaches in the ARG classification task.

Despite the success of these deep learning approaches in the ARG classification task, they all operate on the genome sequence and compare patterns present in that sequence against those of a reference database. However, resistant genes can have dissimilar sequences, but might share similar structures. For instance, the fourth mobile sulfonamide resistance genes have less than 33% similarity to previously known resistance genes, while their protein structures are more than 90% similar to those of known resistance genes [15]. This is because genome evolution accumulates mutations on different bacterial genomes causing dissimilarity in the identified genes. At the same time, natural selection maintains protein structure to produce the same necessary function.

Unfortunately, resolving the structure of a protein is a difficult, time- and cost-intensive task. It requires purifying the protein in the bacterial host and using specialized equipment to determine the tertiary structure through an extensive trial and error approach. This has a severe impact on the number of proteins whose structure has been resolved. More specifically, even though millions of proteins have been sequenced over the past years, only the structures of some thousands of those proteins have been resolved so far. Last years saw a breakthrough in protein structure prediction where the release of the AlphaFold II model [16] enabled the accurate prediction of 3D structures of proteins within only a few minutes.

The success of AlphaFold motivated the development of other approaches designed to predict diverse protein properties. Most of these approaches capitalize on recent advances in natural language processing and in particular on large pre-trained transformer-based language models. These models have been adapted to the setting of proteins and trained in a self-supervised manner on the millions of available protein sequences. Examples of such models include ProtTrans [17] and ProteinBERT [18]. It has been shown that these models go beyond simple pattern matching to perform higher-level reasoning and achieve state-of-the-art performance in many residue-level and protein-level tasks [18, 17, 19]. ESM2 [19] is another recent approach which can be combined with the structure module of AlphaFold [16] and predict the 3D structure of a protein at the resolution of individual atoms with high accuracy.

While these models can develop evolutionary knowledge about proteins, they do not explicitly capture the interactions between amino acids that are in close proximity in the structure space. Therefore, incorporating structural knowledge to these models might potentially lead to performance improvements. This is the main contribution of this paper. More specifically, we propose different methods to classify ARGs based on sequence and/or structure information. The first method explores the power of Large Language Models (LLMs) in classifying the proteins into different antibiotic resistance classes. For that purpose, we fine-tune and evaluate the ESM2 [19] on a standard ARG classification task. The second method is based on a combination of the sequence and the structure of the proteins. First, given a genome sequence, we use the AlphaFold II model to predict its 3D structure. Then, we use both the sequence and the structure to predict the class of the ARG. The sequence is processed by a language model, and the structure by a graph neural network (GNN) [20], which is responsible for capturing the interactions between amino acids that are in close proximity in the space. The second method is also evaluated on the same standard ARG classification dataset. Our results indicate that the proposed methods either outperforms or achieves performance comparable with that of state-of-the-art approaches. Lastly, to study the predictive power of the structure alone, we represented distance matrices (i. e., distance between amino acids in Å) as images and used a CNN to extract relevant features to be used for the classification of proteins. In addition to that, we show that transfer learning can be very helpful in this setting and can improve antibiotic resistance prediction. This is a very important result since the method relies entirely on the structure and has no access to the sequence (e. g., amino acid types, etc.), but still, it managed to yield good performance when CNN models pre-trained on ImageNet [21] are employed.

## Methods

We next give more details about the approaches we developed for the ARG classification problem. More specifically, the first proposed method consists of three main steps: (1) AlphaFold II is first employed to predict the protein structure of the input amino acid sequences; (2) the emerging protein structures are then mapped to graph structures; and (3) the generated graphs are fed into a GNN which interoperates with a large-scale pre-trained protein language model. The second approach differs from the first in that the emerging protein structures are mapped to images and not to graphs, while the generated images are fed into a CNN (instead of a GNN). An overview of the afore-mentioned approaches is illustrated in Figures 1 and 2.

**Figure 1:**
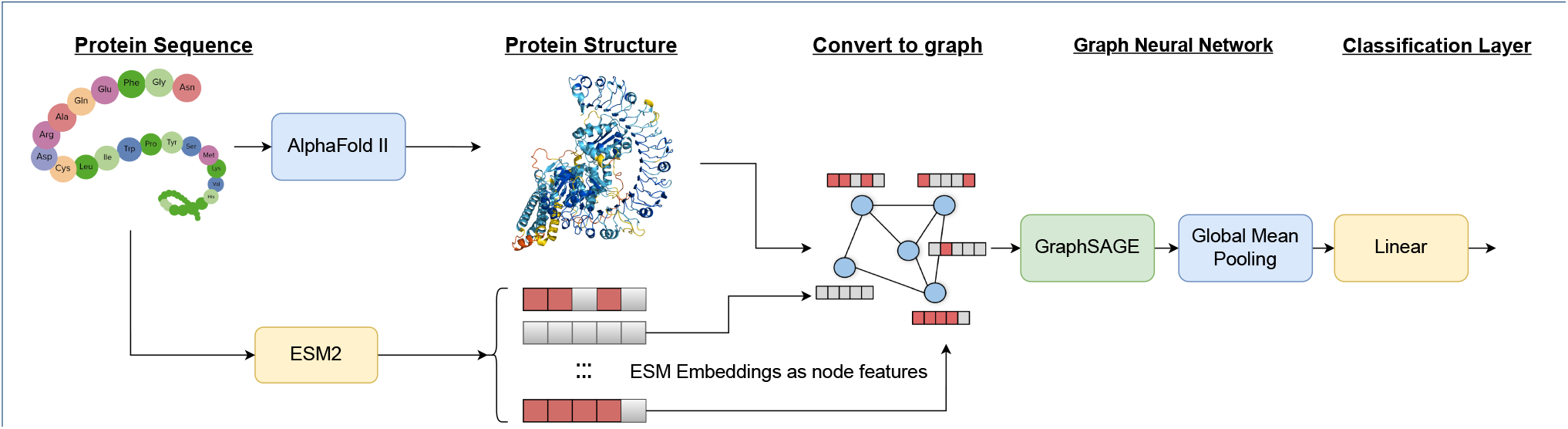
High-level overview of the proposed framework for classifying the different ARGs with GNNs. The framework consists of three main components : (1) The AlphaFold model predicting the 3D structure of an input protein; (2) The protein structures are mapped to graph structures; (3) A classifier, consisting of a GraphSAGE model applied to the graph constructed from the structure and with node features from a pre-trained protein language model (ESM2) applied to the sequence.

**Figure 2:**
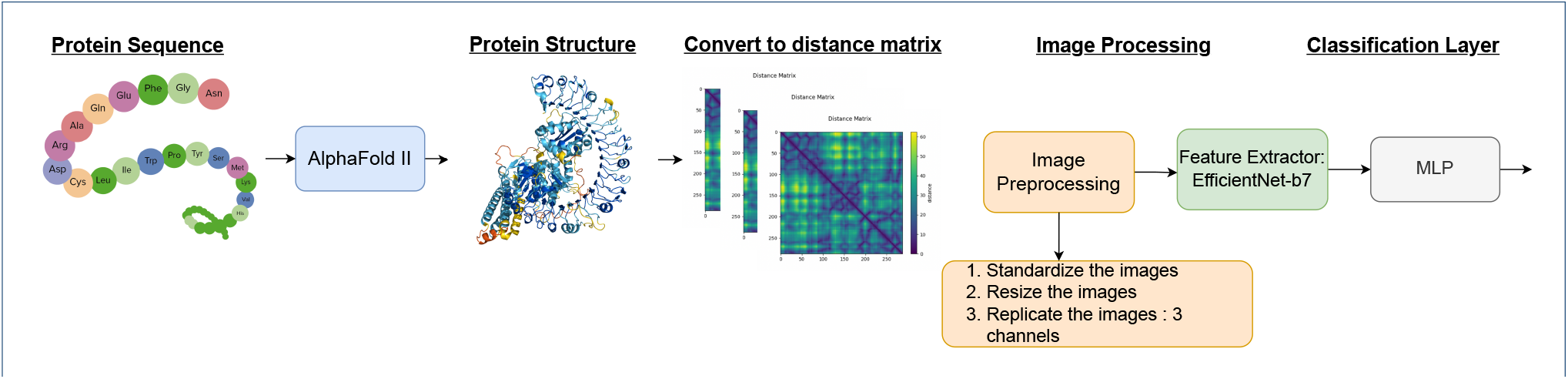
High-level overview of the second approach based on Convolutional Neural Networks. The framework consists of two main components : (1) The AlphaFold model predicting the 3D structure of an input protein; (2) A classifier, consisting of a pre-trained Computer Vision model (EfficientNet-B7) fine-tuned on the classification task of images representing the distance matrices.

### From Sequences to Structures

Note that for the majority of the proteins contained in our dataset, their structure has not been resolved experimentally. Thus, we start by predicting the 3D structure of those proteins. Due to the size of the dataset and limited resources, we made use of ColabFold [22], a software that replaces the MSA generation part of AlphaFold II with an alternative one that is faster and thus reduces the prediction time, which could potentially take several minutes for each protein. One of the key features of AlphaFold II is its ability to refine its predictions through multiple rounds of computation. Even though this process can be time-consuming, from these multiple rounds, AlphaFold II computes five models which help to improve the accuracy of the predictions. There exists a metric (known as pLDDT), which measures the confidence of the system in the predicted structure, with higher scores indicating better predictions. It has been stated that a pLDDT score of 70% or higher is considered to be a threshold above which the predicted structure can be expected to be modelled well [23]. To limit the time required for predictions, we have set a threshold of 70% for the predictions. We report only the first model that meets this threshold with an average per-residue confidence metric.

### From Structures to Graphs

As already discussed, we also use the 3D structure of proteins to enhance the model’s predictive performance. Thus, both sequence and structure are leveraged to determine the resistance class. Following previous studies [24, 25], in this work, we represent the 3D structure as a graph. More specifically, for each protein sequence, AlphaFold II outputs the 3D coordinates of the atoms in the different amino acids. Then, we build a graph *G* = (*V, E*) as follows. Each node *v* ∈ *V* represents an amino acid that exists in the protein, while each edge *e* ∈ *E* connects two amino acids to each other. In our setting, there are two types of edges: (1) peptide bonds which connect residues that are adjacent in the primary sequence; and (2) “proximity” edges that connect (non-bonded) residues which are spatially close to each other. For the second type of edges, we only consider pairs of residues whose distance is less than a pre-defined threshold *ϵ* usually chosen between 6.5 Å and 9.5 Å in the relevant literature [26]. In this work, we set the value of the threshold to 8 Å since it achieves the highest performance on the validation set. Since edges capture distance relationships between nodes, we choose to build a weighted graph. We use the following weighting scheme for the edges:

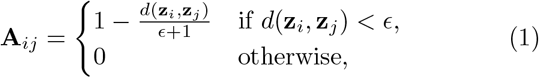

where **z**_*i*_, **z**_*j*_ denote the coordinates of the *C*_*α*_ atoms of the *i*-th and *j*-th amino acid, respectively, *d*(·,·) is the Euclidean distance between vectors, and *ϵ* is equal to the pre-defined threshold of 8 Å discussed above. Note that edge weights typically represent the similarity between the endpoints of the edge, and therefore we transform distances into similarities. We should also mention that a feature vector **x**_*v*_ ϵ ℝ^*d*^ is associated with each node *v* ϵ *V* and more details about the considered features will be given in the forthcoming section.

### From Structures to Images

In this work, we also explore the application of CNNs to images generated from the 3D structure of proteins. We propose a simple approach to turn the structure of a protein into an image. Specifically, we treat matrices that capture the distance between amino acids as images. Formally, for each protein, we compute a matrix **D** ∈ ℝ^*n×n*^ (where *n* is the length of the protein sequence) as follows:

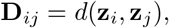

where, once again, **z**_*i*_, **z**_*j*_ denote the coordinates of the *C*_*α*_ atoms of the *i*-th and *j*-th amino acid of the protein, respectively, and *d*(·, ·) is the Euclidean distance between vectors. Note that matrix **D** is sensitive to the ordering of the nodes of the protein (i. e., different orderings give rise to different matrices). However, we can impose an ordering on the nodes (i. e., the ordering of the amino acids in the protein sequence), and thus we can obtain a unique representation for each protein.

Figure 3 illustrates the distribution of the distances between amino acids for 100 randomly sampled proteins from our dataset. To normalize the distances such that they lie in [0, 1], we divide each distance by the average of the maximum values of distance across all proteins. This value was found to be around 130 Å. We then clip the values above the [0, 1] interval (since some of the distances will take larger values than 130 Å). Formally, the values are normalized as follows:

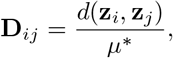

and *µ*^*∗*^ is equal to:

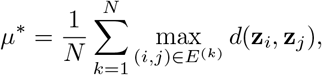

where *N* denotes the number of proteins, 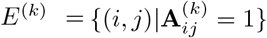 and **A**^(*k*)^ is the adjacency matrix derived from the structure of the *k*-th protein. Then, we multiply the values by 255 and take the integer part, to ensure that all values are integers in {0, …, 255}. CNNs take as input images of fixed size, and therefore, we also resize matrices **D** using bilinear interpolation [27]. Finally, since we employ a CNN model that is pre-trained on ImageNet [21] which requires three channels in the input image (i. e., RGB colors), we replicate the single-channel image into three copies of the same image, representing the different channels. This approach was found to yield better results than modifying the first convolutional layer to take a single channel as input.

**Figure 3:**
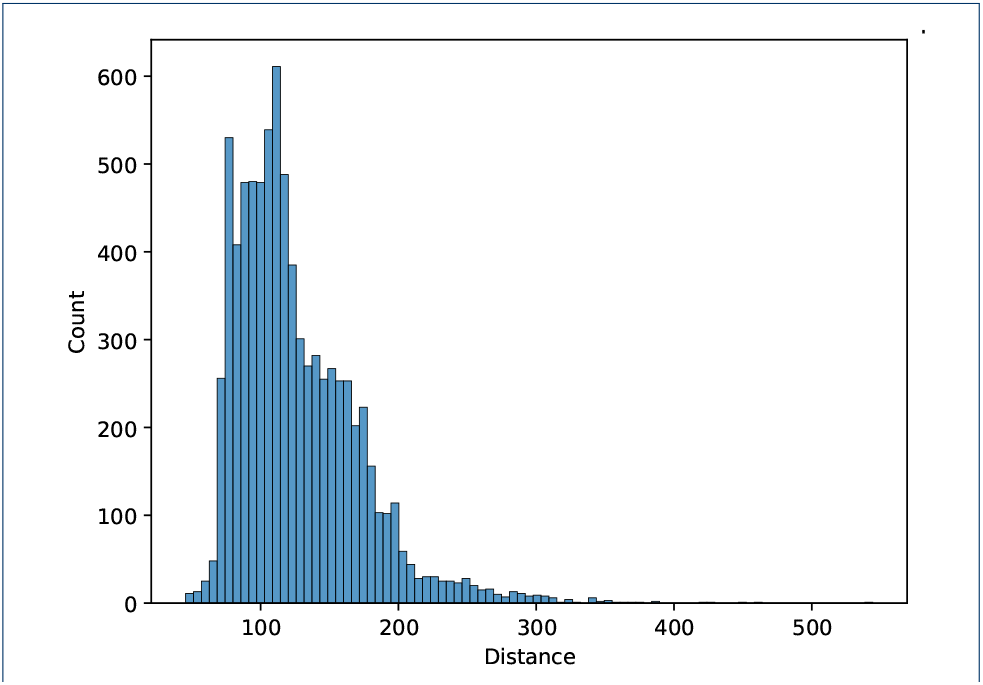
Distribution of the maximum distance between amino acids in each protein from our dataset.

### Classification Models

#### Graph Classification

We employ a message passing GNN to capture the structural information of proteins. The GNN is applied to the graphs that emerge from 3D structures of proteins. To allow the model to learn more complex patterns, we annotate the nodes of the graph with embeddings from a pre-trained language model. Thus, each node *v* ∈ *V* is assigned a feature vector **x**_*v*_ that contains this embedding. To generate those node representations, we employ the Evolutionary Scale Modeling (ESM2) model [19]. ESM2 is the largest pretrained protein language model to date. It mainly consists of transformer layers and was trained on over 60 million protein sequences. It has been shown to be highly performant in predicting the 3D structure of proteins when followed by the structure module of AlphaFold [16], without using multiple sequence alignments. This indicates that the amino acid embeddings it produces are semantically rich and implicitly contain evolutionary information. Note that there are several available variants of ESM2 which usually differ in the number of parameters, ranging from 8 million parameters to 15 billion parameters. Due to limited resources, we only experiment with models that consist of up to 650 million parameters.

While pre-trained language models have shown a high learning ability of the internal representations of proteins and are capable of capturing some functional and structural information, they can still benefit from gaining explicit access to the structure since they can more accurately capture the interactions between amino acids that are in close proximity. To this end, we use the GraphSAGE model [28], a message passing GNN that has been applied with great success to many graph learning tasks. The model consists of a series of neighborhood aggregation layers, where each layer updates the nodes’ feature vectors by aggregating local neighborhood information. Let 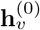 denote the initial feature vector of node *v* which is set equal to **x**_*v*_, i. e., the ESM2 embedding of the amino acid that node *v* represents. In each neighborhood aggregation layer, the hidden state 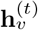 (for *t* > 0) of node *v* is updated as follows:

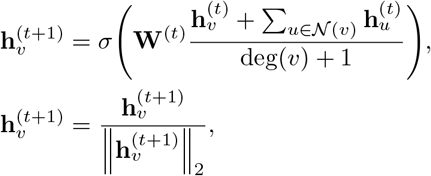

where 𝒩 (*v*) is the set of neighbors of node *v*, deg(*v*) denotes the degree of *v* (i. e., deg(*v*) = |𝒩 (*v*)|), **W**^(*t*)^ is a matrix of trainable parameters and *σ*(·) a non-linear activation function.

Finally, after *T* iterations of neighborhood aggregation, to produce a representation for an entire graph, the model applies the mean operator to the feature vectors of all nodes:

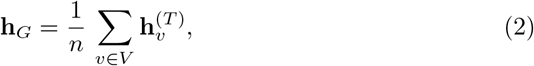

where *n* is the number of nodes of the graph. The graph representation **h**_*G*_ is further fed into a linear layer to produce class probabilities.

#### Image Classification

To classify the image representations of the 3D structure of proteins, we adopt a pre-trained EfficientNetB7 model [29]. The model was trained on the ImageNet dataset, which consists of a diverse range of images. The EfficientNetB7 model employs a compound scaling approach that adjusts the width, depth, and resolution of the network, resulting in a more efficient CNN architecture. The scaling of the depth of the network enables it to capture more complex and sophisticated features. However, this also makes training the network harder due to the vanishing gradient problem [30]. On the other hand, scaling the width of the network allows it to capture finer-grained features and makes it easier to train. Nevertheless, networks that are overly wide and shallow lack the ability to capture high-level features [31]. The use of higher resolution images also allows the CNN to capture finer patterns. To obtain the final class prediction, we added a fully-connected layer that transforms the features extracted by the model into a probability distribution among the different ARG classes. Note that this final layer allows us to perform the ARG classification in an end-to-end manner.

### Implementation

We experimentally evaluate the proposed method in the ARG classification task. We begin by describing the benchmark dataset, the baseline methods and the experimental setup.

#### Dataset

We evaluate the proposed method on the publicly available benchmark dataset “COllection of ALl Antibiotic resistance gene databases”, commonly referred to as COALA [11]. The dataset agglomerates a number of available databases and provides the protein sequences and their corresponding resistance classes. We perform CD-HIT [32] clustering of sequences with a 70% threshold to have a dataset with at most 70% identity between sequences. The identity score between sequences refers to the ratio of identical amino acids in the aligned sequences. We ended up with 11, 090 proteins, each belonging to one out of 16 different classes of antibiotic resistance (see Table 1 for more details). The number of nodes and edges of the generated graphs depend on the considered sequence and threshold *ϵ*. On average, the graphs consist of 281.05 nodes and 1, 624.94 edges, while the average degree of nodes is equal to 5.71. Some classes such as Rifamycin, streptogramin and MLS (macrolide/lincosamide/streptogramin) only contain a small number of training examples which makes the task very challenging.

**Table 1:** Description of the COALA Dataset, along with the splitting. MLS refers to MACROLIDE/LINCOSAMIDE/STREPTOGRAMIN.

| Antibiotic Resistance | Splits 1-5 |  |
| --- | --- | --- |
|  | Train | Test |
| BETA-LACTAM | 3182 | 796 |
| FOLATE-SYNTHESIS-INHABITOR | 1339 | 335 |
| GLYCOPEPTIDE | 1268 | 318 |
| TETRACYCLINE | 813 | 204 |
| AMINOGLYCOSIDE | 771 | 193 |
| TRIMETHOPRIM | 342 | 86 |
| MACROLIDE | 323 | 81 |
| PHENICOL | 264 | 67 |
| QUINOLONE | 157 | 40 |
| SULFONAMIDE | 146 | 37 |
| MULTIDRUG | 118 | 31 |
| FOSFOMYCIN | 53 | 14 |
| BACITRACIN | 45 | 12 |
| MLS | 16 | 5 |
| STREPTOGRAMIN | 14 | 4 |
| RIFAMYCIN | 12 | 4 |

#### Experimental Setup

The different samples were split into training and test sets with an 80 : 20 split ratio with stratification to ensure that class proportions were preserved. The test set was created in such a way that each test sequence has at most 40% identity with any sequence of the training set. To achieve that, we used CD-HIT [32] to define the various clusters. The minimum “in-identity” between sequence pairs within the same cluster is 40%, whereas the maximum “out-identity” between sequence pairs within different clusters is 40%. We iteratively add clusters to the training and test sets, ensuring a stratified split. The whole process was repeated five times giving rise to five splits. No sample appears both in the training set and the test set of a given split. However, the five splits may share some common samples due to the constraint of having at most 40% identity between training and test samples. Some clusters detected by CD-HIT are larger than the size of the test set, and thus can only belong to the training set since, otherwise, the constraint is not respected. For example, each test set has an average of 846 proteins that are totally different of all the other test sets, which represents around 8% of the entire dataset and only 15 proteins are present in all the test sets as indicated by the intersection of the five large ellipses in Figure 4.

**Figure 4:**
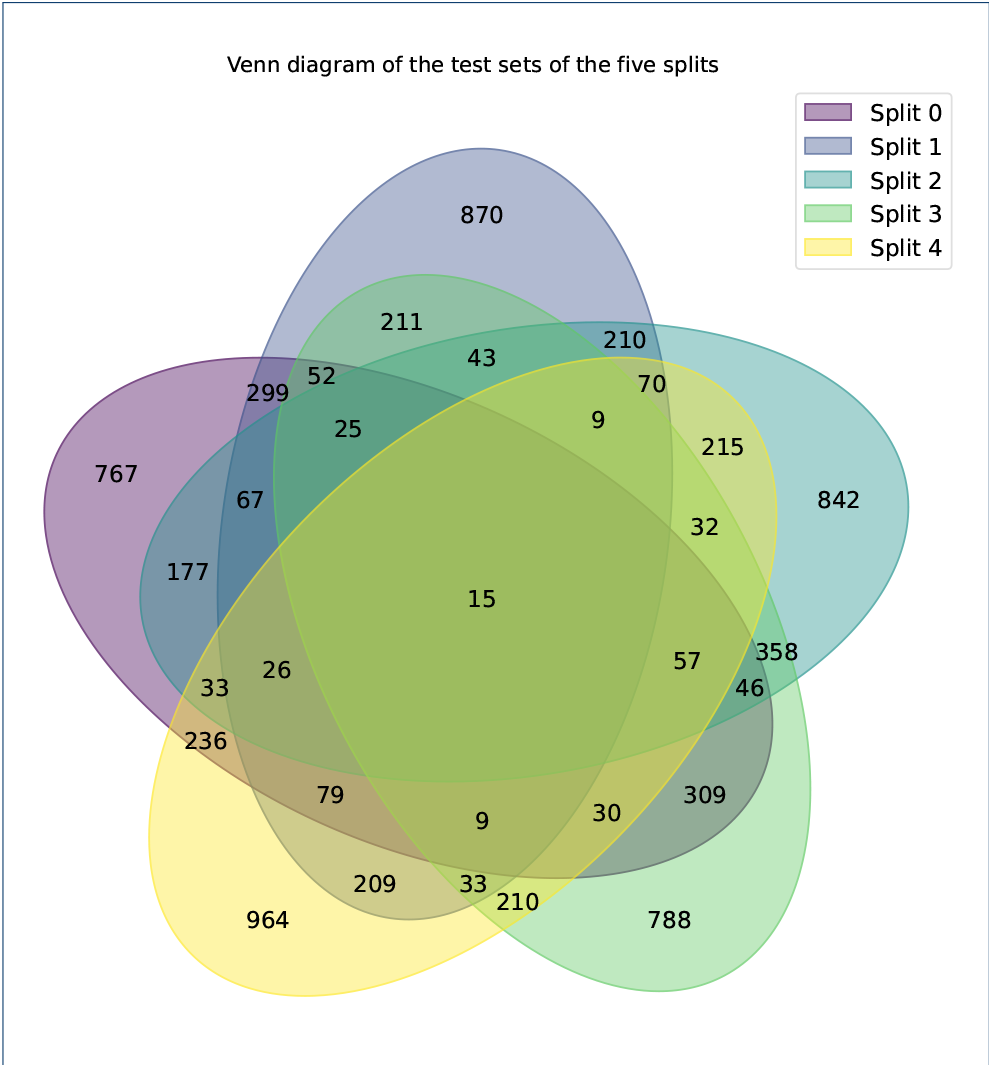
Venn Diagram showing the number of common elements between the five different test sets corresponding to the different data splits.

#### Training and Hyperparameters

We train the full model, consisting of the ESM2 model followed by the GNN model on the COALA dataset, using the Adam optimizer with a learning rate equal to 3 · 10^*−*4^ trying to minimize the cross-entropy loss. We experimented with different message passing GNN architectures (e.g., Graph Convolutional Network [33], Graph Attention Network [34] and GraphSAGE [28]) and we report the results of the best-performing model (GraphSAGE in our setting). We experimented with values of threshold *ϵ* from 5 Å to 15 Å and report the best results (8 Å). We used a dropout rate of 0.2 and a batch size of 32. The GNN model contains 4 hidden layers with a hidden dimension size of 64. We used ReLU activation functions and batch normalization [35] was applied to the output of each GNN layer.

For the CNN-based approach, we also used the Adam optimizer with a learning rate of 3 · 10^*−*4^, trying again to minimize the cross-entropy loss. We set the batch size equal to 8. We experimented with different EfficientNet architectures (from B0 to B7), and reported the best result which in our case emerged from the B7 model. These variants differ in the number of parameters which increases from B0 to B7.

#### Evaluation Metrics

To evaluate the performance of the proposed models, we utilize two metrics: (1) accuracy; and (2) macroaverage F1-score. Accuracy measures the proportion of correct predictions out of all predictions made, while macro F1-score provides a measure of the model’s ability to balance precision and recall across all classes. The definitions of the two metrics are given below:

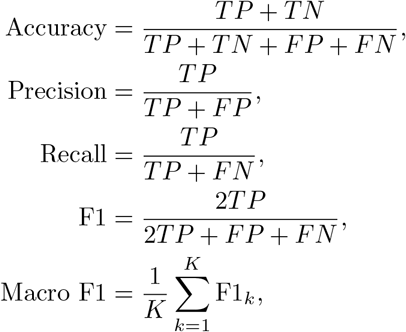

where TP, FP, TN and FN stand for True Positives, False Positives, True Negatives and False Negatives, respectively, *K* is the number of classes and F1_*k*_ is the F1-score of the *k*-th class.

Furthermore, to gain a better understanding of the performance of the model, we also report accuracy with respect to the identity of the best hit in the training set. To do this, we divide the test set into three groups based on the results of a BLAST comparison:

a. The first group consists of elements with a best hit homolog in the training set with an identity greater than 50%.
b. The second group consists of elements with a best hit homolog in the training set with an identity less than 50%.
c. The third group consists of elements with no homolog found in the training set.

This allows us to assess the models’ performance on ARGs that are similar to the training sequences, as well as to evaluate its ability to generalize to novel sequences that are different from the training data.

#### Baselines

Our list of baselines includes mainly homology and sequence-based approaches. Specifically, we compare the proposed method against the following five base-line approaches:

1. BLAST [6] is one of the most powerful tools for sequence alignment. We ran it with max target seqs = 1 and different *e*-value cutoffs (10^*−*3^, 10^*−*7^, 10^*−*10^) as the representatives of the best hit approach.
2. DIAMOND [7] is another tool for sequence alignment. It is faster than BLAST, but less accurate.
3. TF-IDF [36] is a method originally applied in the field of natural language processing to map textual documents to vectors. We generate *n*-gram features for different values of *n* and feed these features to a random forest classification model to make predictions.
4. TRAC [11] employs a transfer learning technique and follows the ULMFit training strategy [12]. The pre-trained TRAC model (LSTM-based language model) was fine-tuned on the COALA dataset with a language model objective, and then fine-tuned for a second time on the classification task.
5. ARG-SHINE [37] is an ensemble method that combines the output of BLAST along with a CNN applied to sequences, and information about the family, domain and motif of proteins. We compare only against their CNN component since it is the dominant component of the model, while the ensemble is based mainly on BLAST results, which can be used with all other models to improve their performance.

## Results & Discussion

### ARG Classification

In Table 2, we report the average prediction accuracy and the corresponding standard deviation across the five folds of the COALA dataset. The two proposed methods (ARG-GNN and ARG-EfficientNet) either outperform or achieve performance comparable with that of state-of-the-art approaches. Specifically, ARG-GNN yields an absolute improvement of 3.10% in accuracy and of 4.08% in macro-average F1-score over TRAC, the best competitor from the baselines. Moreover, the proposed ARG-GNN model performs better than the rest of the methods in the first setting (*<* 50% id) and the third setting, where no homolog is found. Note that BLAST and DIAMOND cannot provide any predictions if there is no homolog found in the training set. Finally, the ARG-GNN model achieves better results than the deep learning methods in the second setting (*>* 50% id), while it is outperformed by both BLAST and DIAMOND. The ARG-EfficientNet model also achieves high levels of performance on the different experimental scenarios of the COALA dataset, but is in all of them outperformed by ARG-GNN. When compared to the baseline methods, ARG-EfficientNet performs relatively well. It out-performs ARG-SHINE and TRAC for sequences with less than 50% identity. However, it yields a lower accuracy than BLAST. For sequences with more than 50% identity, ARG-EfficientNet achieves performance comparable to other deep learning methods. On the other hand, for sequences where no homolog was found, it fails to achieve high levels of performance. To improve the models’ predictive performance even further, we could employ ensemble learning [37], and utilize the output of BLAST when the identity is very high. Overall, the results presented in Table 2 illustrate the superiority of the ARG-GNN model over the baselines, and highlight the potential of deep learning approaches in the task of ARG prediction. Furthermore, the results suggest that the proposed methods can detect ARGs, even when they are not similar to those that belong to the training set. Many novel ARGs contain protein sequences that significantly differ from existing ones, and standard homology-based methods fail to detect them, resulting in a high false negative rate. On the other hand, the proposed models can effectively identify ARGs which are out of the distribution of the training set.

**Table 2:** Classification results (± standard deviation) of the different approaches on the COALA dataset. The best performance in the different groups is typeset in **bold**. N/A refers to “not applicable”.

| Method | Accuracy | Macro F1-score | Accuracy<br>($< 50\%$ id) | Accuracy<br>($> 50\%$ id) | Accuracy<br>(homolog not found) |
| --- | --- | --- | --- | --- | --- |
| BLAST | 65.42% ($\pm 0.57\%$) | 59.29% ($\pm 1.15\%$) | 75.11% ($\pm 1.78\%$) | **95.29%** ($\pm 1.18\%$) | N/A |
| Diamond | 58.86% ($\pm 0.62\%$) | 56.46% ($\pm 2.56\%$) | 64.69% ($\pm 1.72\%$) | 95.11% ($\pm 1.53\%$) | N/A |
| TF-IDF | 54.19% ($\pm 1.62\%$) | 42.18% ($\pm 2.40\%$) | 55.18% ($\pm 1.72\%$) | 84.05% ($\pm 2.07\%$) | 35.43% ($\pm 1.25\%$) |
| TRAC | 69.80% ($\pm 0.66\%$) | 59.70% ($\pm 1.73\%$) | 72.78% ($\pm 1.77\%$) | 90.92% ($\pm 2.16\%$) | 36.83% ($\pm 4.36\%$) |
| ARG-SHINE | 68.34% ($\pm 1.27\%$) | 58.21% ($\pm 2.84\%$) | 69.60% ($\pm 1.78\%$) | 92.88% ($\pm 0.70\%$) | 38.00% ($\pm 3.88\%$) |
| ARG-EfficientNet | 69.83% ($\pm 0.89\%$) | 59.10% ($\pm 2.24\%$) | 73.10% ($\pm 2.17\%$) | 91.67% ($\pm 2.36\%$) | 35.07% ($\pm 2.88\%$) |
| ARG-GNN | **72.90%** ($\pm 0.65\%$) | **63.78%** ($\pm 1.15\%$) | **76.49%** ($\pm 1.30\%$) | 93.00% ($\pm 1.10\%$) | **38.90%** ($\pm 3.98\%$) |

**Table 3:**
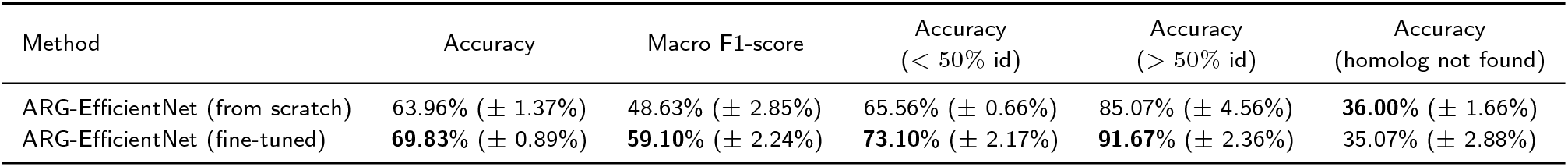
Additional analysis of the effect of transfer learning on the CNN-based classification approach. Classification accuracy (± standard deviation) of the different approaches on the COALA dataset. The best performance in the different groups is typeset in **bold**.

| Method | Accuracy | Macro F1-score | Accuracy<br>($< 50\%$ id) | Accuracy<br>($> 50\%$ id) | Accuracy<br>(homolog not found) |
| --- | --- | --- | --- | --- | --- |
| ARG-EfficientNet (from scratch) | 63.96% ($\pm 1.37\%$) | 48.63% ($\pm 2.85\%$) | 65.56% ($\pm 0.66\%$) | 85.07% ($\pm 4.56\%$) | **36.00%** ($\pm 1.66\%$) |
| ARG-EfficientNet (fine-tuned) | **69.83%** ($\pm 0.89\%$) | **59.10%** ($\pm 2.24\%$) | **73.10%** ($\pm 2.17\%$) | **91.67%** ($\pm 2.36\%$) | 35.07% ($\pm 2.88\%$) |

### Transfer Learning

As discussed above, the proposed ARG-EfficientNet approach achieves performance comparable with that of other deep learning methods in the ARG classification task, but is outperformed by the ARG-GNN model. However, still, this is an interesting approach mainly for the following two reasons:

1. **Sequence-Agnostic**: The model has no access to information about the protein sequence and only limited information about the structure (it has only access to distances of amino acids).
2. **Transfer Learning**: The model was pre-trained on ImageNet, which contains real-world images. These images are vastly different from the “images” constructed from the structure of proteins.

To measure the impact of transfer learning on the performance of the ARG-EfficientNet model, we trained an EfficientNet model from scratch on the ARG dataset. The results are shown in Table 5. We observe that in almost all cases, transfer learning leads to significant improvements in performance. For instance, pre-training on ImageNet offers an absolute improvement of 5.87% in accuracy and of 10.47% in macro-average F1-score. Transfer learning only fails to yield improvements in the case of sequences where no homolog was found. The obtained results highlight the power of transfer learning, where a pre-trained model can perform significantly better than a model trained from scratch, even on a different domain.

### Impact of Sequence and Structure

We next explore what is the impact of information extracted from the protein sequence and structure on the performance of the proposed ARG-GNN model. More specifically, we conducted the following ablation study. First, a GraphSAGE model (i. e., the backbone of ARG-GNN) was trained and evaluated on the COALA dataset, for the task of predicting resistance. The nodes of the graph representations of proteins were annotated with the following features: the vectorized representations (i. e., one-hot encoding) of amino acid types, the *ϕ* and *ψ* torsion angles, and also the relative solvent accessibility (RSA) of each amino acid which reflects its degree of exposure or concealment in the 3D structure. This model hence has access to information from the protein structure only. We also performed another experiment where we fine-tuned the ESM Language model on the classification task. We fed the BOS (Beginning of Sequence) token embedding to a multi layer perceptron (MLP) to produce the output. This model hence has access to information extracted from the sequence only. The results of these experiments are reported in Table 4. We observe that ESM outperforms GraphSAGE under all scenarios. We hypothesize that this is because ESM has been pre-trained on a very large number of proteins. We also observe that the proposed ARG-GNN approach (i. e., combination of ESM and GraphSAGE) shows superior performance compared to its two components, providing evidence for the hypothesis that the infusion of structural information into sequence-based methods facilitates the learning process and enhances their predictive capability.

**Table 4:** Additional analysis of the utility of combing GraphSAGE and the ESM model. Classification accuracy (± standard deviation) of the different approaches on the COALA dataset. The best performance in the different groups is typeset in **bold**.

| Method | Accuracy | Accuracy<br>($< 50\%$ id) | Accuracy<br>($> 50\%$ id) | Accuracy<br>(homolog not found) |
| --- | --- | --- | --- | --- |
| ESM | 70.86% ($\pm 1.33\%$) | 74.37% ($\pm 1.70\%$) | 92.60% ($\pm 0.50\%$) | 35.45% ($\pm 4.93\%$) |
| GraphSAGE | 66.67% ($\pm 0.98\%$) | 69.38% ($\pm 1.68\%$) | 89.72% ($\pm 1.48\%$) | 32.57% ($\pm 3.18\%$) |
| ARG-GNN | **72.90%** ($\pm 0.65\%$) | **76.49%** ($\pm 1.30\%$) | **93.00%** ($\pm 1.10\%$) | **38.90%** ($\pm 3.98\%$) |

**Table 5:** Additional analysis of the effect of ESM2 size and the effect of injecting structural bias into the different ESM2 variants. Classification accuracy (± standard deviation) of the different approaches on the COALA dataset. The best performance in the different groups is typeset in **bold**. Inf. Time refers to the inference time of the model on one NVIDIA Quadro RTX 6000/8000 gpu with batch size of 4. In number of parameters (Nb. Params), M refers to million and K to thousand.

| Method | Nb. params | Inf. time (seconds) | Accuracy | Accuracy (< 50% id) | Accuracy (> 50% id) | Accuracy (homolog not found) |
| --- | --- | --- | --- | --- | --- | --- |
| ESM 8M. | 7.5M | 5 | 70.86% (± 1.33%) | 74.37% (± 1.70%) | 92.60% (± 0.50%) | 35.45% (± 4.93%) |
| ESM 35M. | 33.5M | 10 | 71.35% (± 0.89%) | 75.1% (± 1.56%) | 92.45% (± 0.71%) | 35.74% (± 2.98%) |
| ESM 150M. | 148M | 25 | 71.58% (± 1.39%) | 74.91% (± 2.17%) | 92.42% (± 1.25%) | 37.74% (± 4.31%) |
| ESM 650M. | 651M | 55 | **73.11%** (± 1.38%) | **76.52%** (± 1.74%) | **90.94%** (± 0.80%) | **42.16%** (± 3.57%) |
| ARG-GNN (ESM 8M.) | 7.5M+785K | 7 | **72.90%** (± 0.65%) | **76.49%** (± 1.30%) | 93.00% (± 1.10%) | **38.90%** (± 3.98%) |
| ARG-GNN (ESM 35M.) | 33.47M+785K | 12 | 71.42% (± 1.17%) | 74.48% (± 1.69%) | 92.79% (± 0.58%) | 37.92% (± 4.41%) |
| ARG-GNN (ESM 150M.) | 147M+785K | 28 | 70.40% (± 1.18%) | 73.59% (± 2.19%) | 91.35% (± 1.61%) | 36.91% (± 2.64%) |
| ARG-GNN (ESM 650M.) | 649M+785K | 65 | 72.27% (± 1.03%) | 75.46% (± 1.60%) | **93.25%** (± 1.00%) | 38.76% (± 2.05%) |

### Other ESM Variants

It should be mentioned that the above findings are based on the ESM2 variant which consists of 8 million parameters, and which is the smallest of all available model variants. It was created as part of research to investigate the influence of scaling language models on their structure prediction performance, and the largest models perform significantly better. We also experimented with other variants of the ESM2 model that consist of different number of parameters. The results, provided in Table 6, show that the larger the ESM2 models (in terms of the number of parameters), the higher its performance in the task of ARG classification. However, when integrated into the ARG-GNN model, it was observed that the larger models did not yield any improvements in performance in most cases. For instance, the ESM2 model that consists of 650 million parameters is the best-performing method only in one experimental scenario. We attribute this phenomenon to the presence of structural information in the larger model’s embeddings. An appealing property of the proposed ARG-GNN model is that it does not require extensive computational resources when structural information is already available. In fact, when combining the ESM2 model that contains 8 million parameters with a GraphSAGE model of approximately 785 thousand parameters, performance was on par with the largest ESM2 model while being 80 times smaller in size. In addition to that, it is also much faster in inference time (7 seconds for the ARG-GNN (ESM2 8M) model vs. 55 seconds for the ESM 650M model).

**Table 6:**
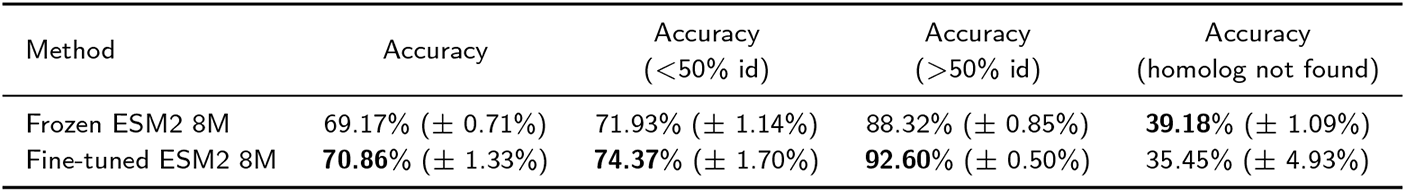
Additional analysis of the effect of finetuning ESM2. Classification accuracy (± standard deviation) of the different approaches on the COALA dataset. The best performance in the different groups is typeset in **bold**.

We also evaluated the performance of the raw pretrained ESM2 embeddings in the considered resistance prediction task. The results given in Table 6 show that the raw (i. e., frozen) ESM2 embeddings achieve similar levels of performance in most cases to the fine-tuned ones. This highlights the fact that the ESM2 embeddings capture different properties of proteins and can thus be directly utilized in downstream tasks without the need for additional fine-tuning on task-specific data.

### Limitations

We next discuss some limitations of the proposed method, which leverages both the sequence and structure of proteins to deal with downstream tasks. First, the proposed approach depends significantly on the availability of the structure of proteins. Even though the structures of a very large number of proteins have been resolved so far, there still exist proteins for which the structure is not known. Furthermore, predicting the structure of a protein using complex modeling techniques such as AlphaFold II can be both a time-consuming and resource-intensive procedure. In addition, the deployment of LLMs, despite their demonstrated success in similar applications, entails substantial computational overhead due to the large number of parameters. We should also note that the performance of the proposed approach is contingent upon the quality and quantity of training data. Although our proof-of-concept results are promising, however, for practical applications, a larger and more diverse labeled dataset is necessary, particularly to ensure adequate representation of each class.

## Conclusions

This work investigates how deep learning approaches can be utilized to detect antibiotic resistance by incorporating information from both protein sequence and also from structure. Our results show that the integration of structural bias into pre-trained language models on protein sequences leads to improved performance compared to the state-of-the-art methods in the case of smaller language models. However, no significant improvement was observed for larger language models potentially due to the fact that they incorporate information about the structure of proteins. Our results highlight the advantage of using smaller protein language models when the structure is available, as they are computationally more efficient and in our setting as effective as larger models. Quite surprisingly, we also found that the fine-tuned EfficientNetB7 model, which is applied to the structure solely, performed well, demonstrating also the potential of transfer learning in this domain.

This work could be of interest to people from different backgrounds. For computer scientists, it offers a new way to combine sequence and structural information and showcases the impact of transfer learning. For biologists, it highlights the potential of deep learning in the field and provides a model that can be applied to any labeled datasets to make predictions and explanations. In terms of future research, we plan to explore methods for integrating supplementary information sources, such as functional annotations, into the model. Furthermore, combining multiple language and graph neural network models into an ensemble could potentially enhance the performance of the proposed approach.

## Acknowledgements

We are grateful to Dr. A. Tsortos for useful discussions during the initial stages of this project.

## Availability of data and materials

The original COALA dataset is available at: https://figshare.com/articles/dataset/COALA_datasets/11413302. The source code for training our model together with the pre-processed dataset is available at https://bit.ly/3K6AYLz.

## Ethics approval and consent to participate

Not applicable.

## Consent for publication

Not applicable.

## Authors’ contributions

A.Q. and S.E. initiated the study. A.Q. and S.E. developed the methodology under the supervision of M.V. while A.Q was the main implementer of the deep learning models. A.Q., S.E. and G.N. wrote the manuscript. All authors were involved in the discussion and finalization of the manuscript. All authors reviewed the manuscript before submission for publication.

